# Frequent Monitoring of C-peptide Levels in Newly Diagnosed Type 1 Subjects Using Dried Blood Spots Collected at Home

**DOI:** 10.1101/286237

**Authors:** Ruben H. Willemsen, Keith Burling, Peter Barker, Fran Ackland, Renuka P. Dias, Julie Edge, Anne Smith, John Todd, Boryana Lopez, Adrian P. Mander, Catherine Guy, David B. Dunger

## Abstract

**Objective:** To evaluate a novel approach to measure ß-cell function by frequent testing of C-peptide concentrations in ‘dried blood spots’ (DBS)

**Patients:** Thirty-two children, aged 7-17 years, recently diagnosed with type 1 diabetes.

**Design:** Mixed-meal-tolerance-test (MMTT) within 6 and again 12 months after diagnosis with paired venous and DBS C-peptide sampling at 0 and 90 minutes. Weekly DBS C-peptide before and after standardized breakfasts collected at home.

**Results:** DBS and plasma C-peptide levels (n=115) correlated strongly (r=0·91; p<0.001). The Bland-Altman plot indicated good agreement. The median number of home-collected DBS cards per participant was 24 over a median of 6.9 months. Repeated DBS C-peptide levels varied considerably within and between subjects. Adjustment for corresponding home glucose measurements reduced the variance permitting accurate description of changes over time. The correlation of the C-peptide slope over time assessed by repeated home DBS versus area under the curve during the two MMTTs was r=0·73; p<0.001. Mixed models showed that a 1-month increase of diabetes duration was associated with 17 pmol/l decline in fasting DBS C-peptide, whereas increases of 1 mmol/l in glucose, 1 year older age-at-diagnosis and 100 pmol/l higher baseline plasma C-peptide were associated with 18, 17 and 61 pmol/l higher fasting DBS C-peptide levels, respectively. In addition, glucose responsiveness decreased with longer diabetes duration.

**Conclusion:** Our approach permitted frequent assessment of C-peptide, making it feasible to monitor ß-cell function at home. Evaluation of changes in the slope of C-peptide using this method may permit short-term evaluation of promising interventions.

## Introduction

Our knowledge of the natural history of type 1 diabetes (T1D) has changed over the last few years with the increasing awareness that a period of dysglycaemia and reduced ß-cell glucose sensitivity may occur between the first appearance of autoantibodies and the development of disease (1–3). Even when T1D has developed the rate of immune-mediated destruction of ß-cells may vary considerably with some subjects becoming C-peptide deplete soon after presentation and others maintaining some degree of ß-cell function for many years after diagnosis (4–8). Thus, our ability to monitor changes in ß-cell function over the short and longer term may be critical to our understanding of disease pathophysiology and also the evaluation of potentially useful interventions to prevent the development and progression of the disease (9).

C-peptide provides an excellent marker of residual ß-cell activity and even random or fasting levels may be of clinical significance as they have been associated in longitudinal studies with HbA1c, risk for microvascular complications and the incidence of hypoglycaemia (8,10-12). Little is known about the day-to-day variation in random or fasting C-peptide levels in people who developed T1D or who are in that pre-diabetic period. It has been assumed that such data are likely to be highly variable and that more formal assessments of residual ß-cell function are required for the evaluation of therapeutic interventions. The mixed meal tolerance test (MMTT) and glucagon stimulation tests have been shown to provide a good measure of ß-cell function (13) and in an FDA issued draft guidance it was suggested that the repeated MMTT might be the best standardised measure of drug efficacy in Phase 2 and 3 clinical trials of potentially useful interventions (14). A basic protocol based on repeated MMTT over 1-3 years has remained the benchmark for all subsequent clinical studies in pre-diabetes and newly diagnosed T1D.

The MMTT however, is quite labour intensive and requires admission to a clinical research facility for several hours. It is therefore probably not the best test to obtain repeated measures of C-peptide over relatively short periods of time. There is increasing interest in determining how environmental or short-term drug exposures affect ß-cell function. To develop methods that might be suitable for this purpose we have examined the value of fasting and single post-prandial measures of C-peptide collected using dried blood spots in the home setting. We report longitudinal changes in fasting and post-prandial C-peptide in young people recently diagnosed with T1D and contrast the information gained about change in C-peptide over time with that obtained by more formal testing of interval MMTT.

## Subjects and Methods

### Subjects

Thirty-two subjects with T1D, aged 5-18 years, with ≥1 auto-antibody positive in their local laboratory, requiring insulin treatment, were recruited within 1-24 weeks from diagnosis.

Exclusion criteria: type 2, monogenic, and secondary diabetes, use of immunosuppressive agents or oral steroids in the last two months, pregnancy and coeliac disease. Four participants withdrew before, and two withdrew at the second MMTT (one unable to cannulate, one unable to perform MMTT owing to hyperglycaemia and not wanting to be rescheduled).

### Study design

The patients underwent two MMTTs (Figure 1): the first one at recruitment (any time between 1-24 weeks from diagnosis) (visit 1), and the second 12 months from diagnosis (visit 2) (Figure 1). In between, they were asked to collect weekly DBS at home before (fasting) and 90 minutes after a standardised breakfast, with paired recordings of capillary glucose on their own glucometer. Written and verbal instructions for DBS collection were provided at visit 1. Participants were instructed to let the DBS card dry for 24 hours without using heat or direct sunlight, and to post the cards to the laboratory via pre-paid envelopes. Participants were provided with a list of standardised breakfasts containing 35, 45 and 50 grams of carbohydrates for 5-6, 7-12 and 13-18 year olds respectively, and were asked to pick a breakfast which could be eaten on a weekly basis.

**Figure 1.**
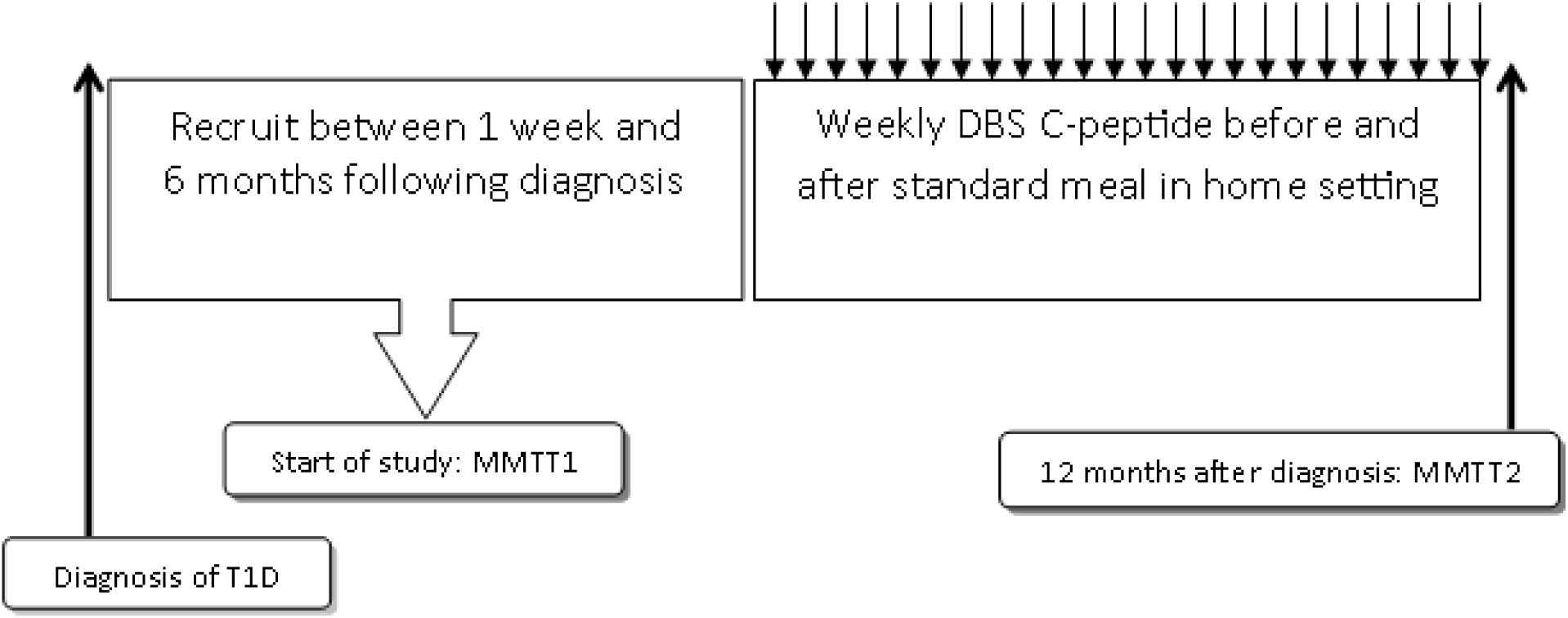
Study design

### Mixed Meal Tolerance Test

The MMTT was performed following an overnight fast (from midnight), with no food or drink other than water. Long-acting insulin and basal rates (for insulin pump users) were continued as normal. The use of rapid-acting insulin was acceptable up to two hours before the MMTT. The MMTT was only performed if the participant’s glucose level was 4-11.1 mmol/l. Participants ingested 6 ml/kg of Boost meal solution (maximum 360 ml; Nestlé HealthCare Nutrition), within 10 minutes. Blood samples for C-peptide and glucose were collected 10 minutes prior to the meal (-10 min), at time of ingestion (0 minutes), and at 15, 30, 60, 90 and 120 minutes.

Two capillary DBS samples were taken during the MMTT: between -10 and 0 minutes, and at 90 minutes for comparison with plasma samples.

### Other assessments

At each MMTT, height and weight were measured and BMI was calculated. Age- and sex-appropriate standard deviation scores were calculated for height, weight and BMI (15, 16).

HbA1c, insulin regime and total daily dose of insulin/kg ((mean insulin requirements over the last 3 days)/weight) were recorded at both MMTT visits.

### Laboratory methods

Plasma C-peptide samples taken during the MMTT were assayed in singleton on a DiaSorin Liaison® XL automated immunoassay analyser using a one-step chemiluminescence immunoassay. Between-batch imprecision for the assay is 6·0% at 561 pmol/L (n=181), 4·3% at 2473 pmol/L (n=170), & 5·4% at 5400 pmol/L (n=166).

DBS cards were stored at -80° C until analysis. DBS C-peptide was analysed in duplicate using a modification of the Mesoscale Discovery two-step electrochemical immunoassay (MSD Custom human C-peptide kit). Two 3·2mm diameter spots were punched for each standard, quality control and sample and eluted into MSD Diluent 13 by shaking for one hour at 4° C. A portion of this elution buffer was transferred to the analytical plate and the standard two-step immunoassay was performed.

For the DBS standards EDTA blood was spun and plasma removed. Red cells were washed three times in 1x Phosphate Buffered Saline (PBS) to remove endogenous C-peptide. The red cell pellet was partially reconstituted in 5% Bovine Serum Albumin (BSA) in 1xPBS. The C-peptide IRR was prepared in 5% BSA in 1xPBS. The prepared standards were diluted 1:4 in the red cell prep to produce DBS standards ranging from approximately 4000 to 125 pmol/l, based on the serum concentration, in artificial whole blood with a packed cell volume of approximately 0.45. This artificial whole blood was spotted out, allowed to dry and stored - 80°C. The assigned value was derived by triplicate analysis of the plasma from the whole blood standards using the DiaSorin Liaison assay.

Quality control data analysed over a seven-month period showed no noticeable trend in results, indicating good stability over time. Between-batch imprecision for the DBS assay was 8·7% at 451 pmol/L (n=115), 10·0% at 495 pmol/L (n=115), and 11·3% at 878 pmol/L (n=113). The lower limit of detection (LLD) was 50 pmol/l. Values recorded as lower than LLD were assigned 25 pmol/l.

Glucose levels taken at the MMTT were analysed via an adaption of the hexokinase-glucose-6-phosphate dehydrogenase method (17). For the home glucose measurements, patients used a range of glucometers.

### Ethics

The National Research Ethics Committee East of England – Cambridge South approved the study. All patients ≥16 years and parents gave informed consent, and children <16 years gave assent to the study procedures.

### Statistical analyses

Data are expressed as mean (SD) or median (IQR), depending on normality of data distribution. Changes in parameters between visit 1 and 2 were evaluated by a paired Student’s t-test or Wilcoxson’s paired rank test.

Intra-class correlation was used to compare the paired plasma and DBS samples. A power calculation showed that, based on a two-sided test, a sample size of 30, allowing for a drop-out rate of 20%, alpha 5%, provided >90 % power to detect a correlation coefficient of 0·6 or higher between plasma and DBS C-peptide levels. In addition, a Bland-Altman plot (the differences between DBS and plasma C-peptide plotted against the mean of those measurements) was produced to compare the plasma and DBS measurements. As the differences between DBS and plasma C-peptide were not normally distributed, data were log transformed prior to calculating the mean and the difference.

For the correlation analysis and Bland-Altman plot all available paired samples were used (n=115; three missing DBS samples). For the analyses of post-prandial DBS C-peptide and DBS C-peptide increment, only the home recordings were used, as the samples collected during the MMTT visits, were collected after a different stimulus and without insulin administration.

To compare changes in ß-cell function across the two methods (AUC C-peptide during MMTT versus post-prandial DBS C-peptide) slopes were estimated by regressing each estimate of ß-cell function on duration of diabetes. The estimated slopes were then divided by their respective standard deviation to standardise the measurements in scale. This produces a measurement akin to a Z-score, since the measurements have been standardised in scale but not centred at the mean. Bland-Altman plots were produced to compare the two methods.

### Correlation was calculated using Pearson correlation

To investigate the effects of diabetes duration, glucose levels, baseline plasma C-peptide, age at diagnosis and sex on DBS C-peptide, mixed effects regression models were used with random intercepts to account for the repeat measurements per person. In these models diabetes duration, glucose levels, baseline plasma C-peptide and age at diagnosis were entered as covariates and sex as a factor. To examine whether there was a change in glucose responsiveness with longer duration of diabetes, we tested whether there was an interaction between diabetes duration and glucose levels. To examine whether there was a more pronounced decline in C-peptide with longer diabetes duration in younger children, we tested whether there was an interaction between age at diagnosis and diabetes duration.

Statistical analysis was performed in SPSS version 23 and R version 1.0.136.

## Results

### Baseline data

Table 1 shows parameters of all participants at visit 1 and 2. HbA1c deteriorated slightly over time. The insulin dose did not change significantly. Fasting and 90 min glucose levels increased significantly. As expected, all estimates of ß-cell function (DBS C-peptide, AUC and peak plasma C-peptide derived from the MMTT) decreased significantly over time.

**Table 1.** Baseline data

|  | **Visit 1** | **Visit 2** | **P-value** |
| --- | --- | --- | --- |
| N | 32 | 28 | - |
| Ethnicity | Asian 3<br>Mixed 3<br>White 26 | Asian 3<br>Mixed 3<br>White 22 | - |
| Age at diagnosis (yr) | 12.0 (2.9) | 11.9 (3.1) | - |
| Age at visit (yr) | 12.4 (2.9) | 12.9 (3.1) | p<0.001 |
| Sex (m/f) | 12/20 | 9/19 | - |
| Duration of diabetes (days)* | 154 (65) | 365 (9) | p<0.001 |
| HbA1c (mmol/mol) | 51 (12) | 59 (20) | p<0.05 |
| Height SDS | 0.19 (1.0) | 0.15 (1.0) | NS |
| BMI SDS | 0.62 (1.0) | 0.64 (1.2) | NS |
| Insulin regimen | Basal bolus 27<br>Pump 5<br>3 or less injections<br>0 | Basal bolus 18<br>Pump 9<br>3 or less injections<br>1 | - |
| Insulin dose (U/kg/day) | 0.57 (0.23) | 0.64 (0.28) | NS |
| Fasting glucose (mmol/l)* | 6.5 (2.3) | 8.1 (2.8) | p<0.01 |
| Fasting DBS C-peptide (pmol/l)* | 379 (396) | 245 (240) | p<0.05 |
| 90 min glucose (mmol/l)* | 15.4 (4.0) | 19.0 (3.9) | p<0.001 |
| 90 min DBS C-peptide (pmol/l)* | 948 (727) | 493 (641) | p<0.001 |
| Mean AUC C-peptide (pmol/l)* | 0.59 (1.76) | 0.34 (0.45) | p<0.001 |
| Peak C-peptide in MMTT (pmol/l)* | 794 (582) | 422 (660) | p<0.001 |

All values reported as mean (SD), except for \* (median (IQR)).

### Comparison of the plasma and DBS C-peptide method

DBS and plasma C-peptide levels correlated strongly (n=115 paired samples; r=0·91; p<0·001; Figure 2). The Bland-Altman plot is shown in Figure 3. The DBS method slightly over-estimated C-peptide levels with a mean (SD) difference of 1·27 (1·31) times the plasma values (p<0·001). 95% limits of agreement were 0·76-2·18.

**Figure 2.**
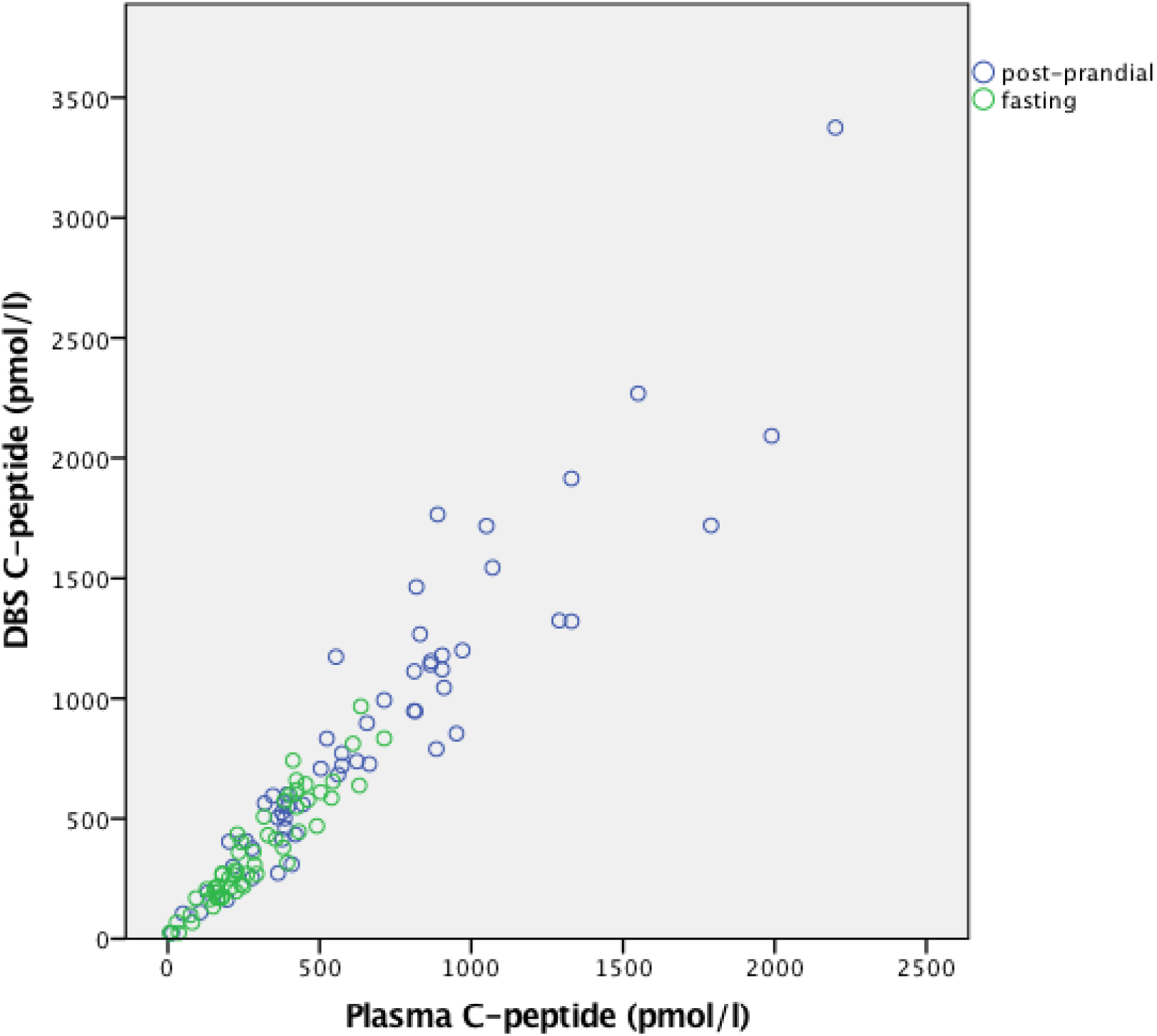
Correlation between plasma and DBS C-peptide levels. Fasting and post-prandial values indicated by green and blue circles, respectively. N=115; r=0·91; p<0·001

**Figure 3.**
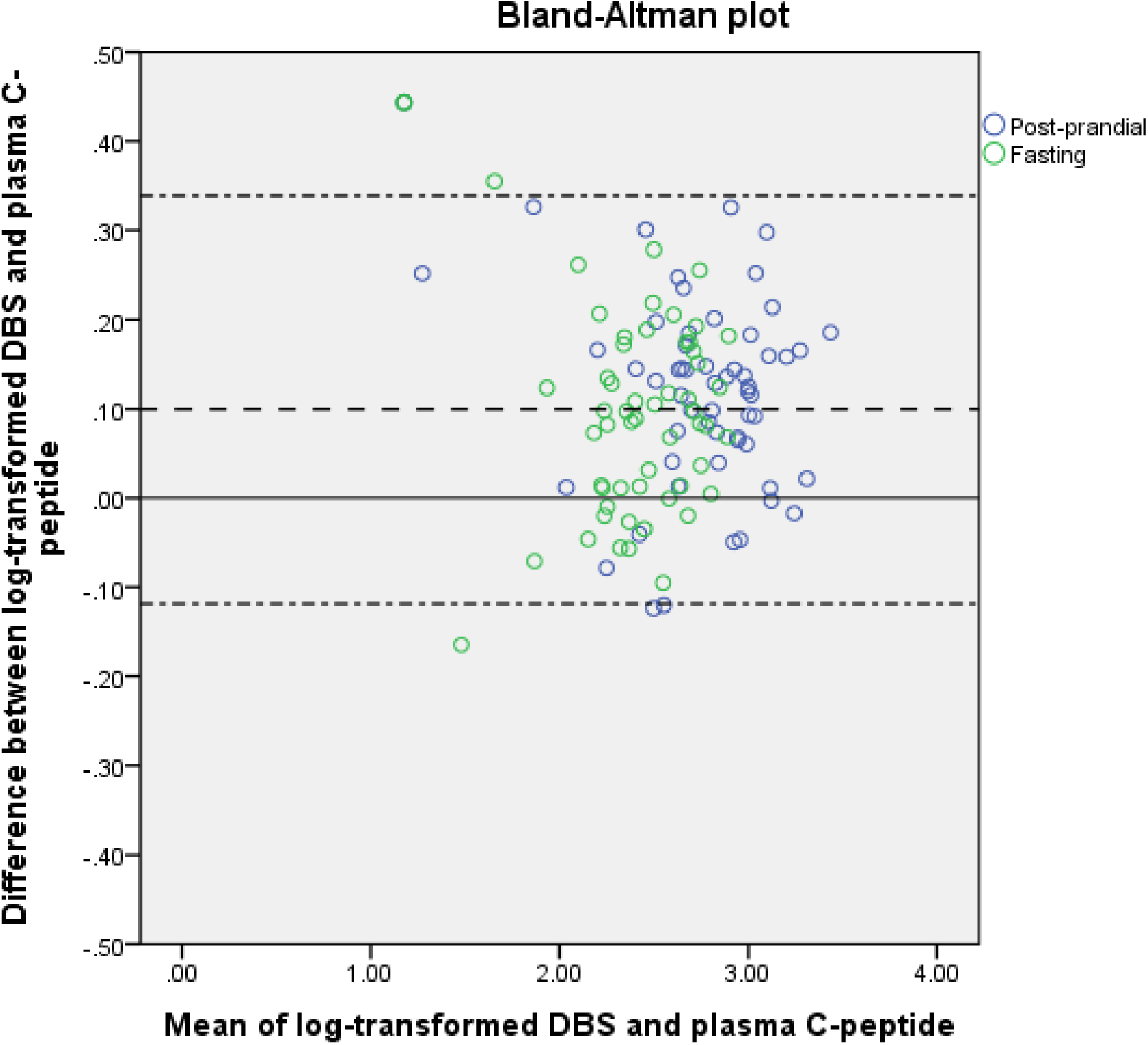
Bland-Altman plot for the comparison of the DBS and plasma C-peptide assay

### DBS C-peptide measurements

For those that completed the study up to the second MMTT, the median number of home DBS samples per participant was 24 (min. 8 and max. 40), collected over a median duration of 6.9 months (IQR 1.8). All but two participants had detectable C-peptide levels throughout the study. Supplementary figure 1 shows the course of fasting and post-prandial DBS C-peptide measurements over time from each subject.

Comparison of slopes defined by the different methods to estimate the change in ß-cell function over time

We compared the change in ß-cell function over time as estimated by the AUC plasma C-peptide during the MMTT (‘MMTT’) versus post-prandial home DBS C-peptide (‘DBS’) respectively, by constructing a Bland-Altman plot where the slopes derived from the two methods were compared (Figure 4). The mean (SD) difference in slopes was 0·10 (0·75) for DBS versus MMTT (not significantly different from zero). The correlation of the slopes between the two methods (DBS versus MMTT) was r=0·73, p<0·001.

**Figure 4.**
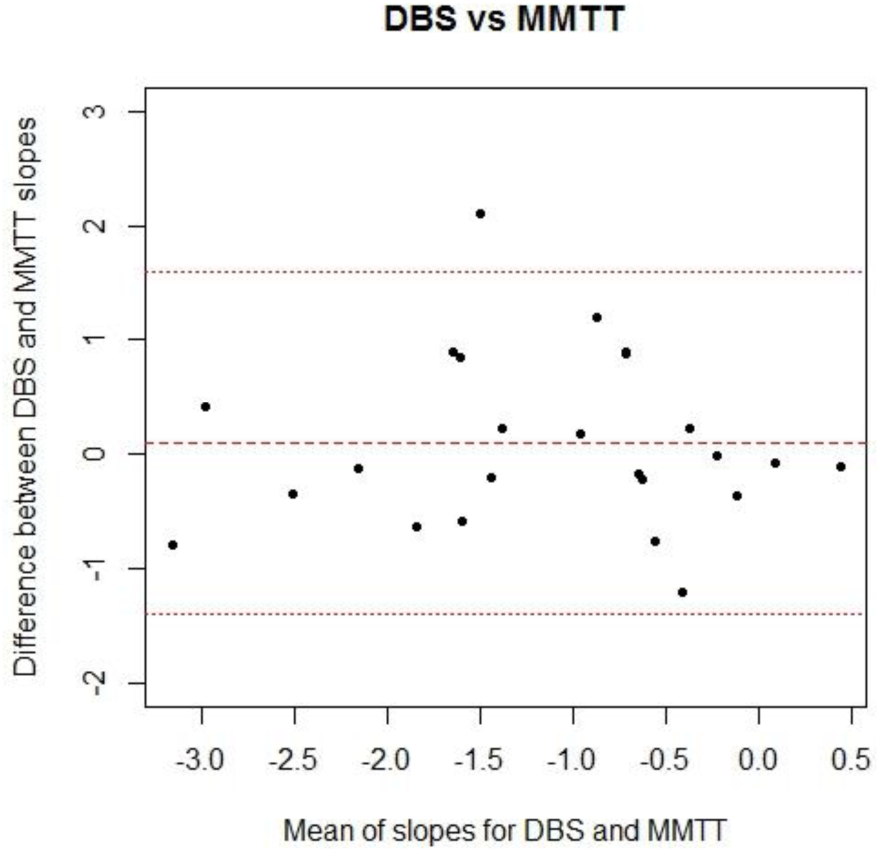
Bland-Altman plot for the comparison of the slopes in ß-cell function across the 2 different methods

### Covariates of fasting DBS C-peptide

In a series of models we evaluated covariates of fasting DBS C-peptide (Table 2A). Diabetes duration had a significant negative effect on fasting DBS C-peptide. Fasting glucose, age at diagnosis and baseline fasting C-peptide had significant positive effects on subsequent DBS C-peptide levels (Models 1 and 2). Sex did not have a significant effect (data not shown).

**Table 2A.** Mixed model analysis of variance: fasting DBS C-peptide

|  | **Model 1** |  | **Model 2** |  | **Model 3** |  | **Model 4** |  |
| --- | --- | --- | --- | --- | --- | --- | --- | --- |
|  | Beta | p-value | Beta | p-value | Beta | p-value | Beta | p-value |
| Diabetes duration (days) | -0.57 | <0.001 | -0.57 | <0.001 | -1.2 | <0.01 | -1.8 | <0.001 |
| Fasting glucose (mmol/l) | 17.6 | <0.001 | 17.9 | <0.001 | 45.1 | <0.001 | 43.5 | <0.001 |
| Age at diagnosis (years) | 25.7 | 0.025 | 17.2 | 0.037 | 16.8 | 0.044 | 7.4 | 0.433 |
| Baseline plasma C-peptide (pmol/l) |  |  | 0.61 | <0.001 | 0.60 | <0.001 | 0.60 | <0.001 |
| Diabetes duration x fasting glucose<sup>§</sup> |  |  |  |  | -0.10 | <0.001 | -0.10 | <0.001 |
| Diabetes duration x diabetes duration (days<sup>2</sup>)<sup>§</sup> |  |  |  |  | 0.003 | <0.001 | 0.003 | <0.001 |
| Diabetes duration x age at diagnosis<sup>§</sup> |  |  |  |  |  |  | 0.04 | 0.051 |
| **Akaike's Information Criteria** | 8941 |  | 8920 |  | 8903 |  | 8901 |  |

Consequently, we investigated whether glucose responsiveness changed over time by adding an interaction term (Model 3), and this further improved the model, indicating that the C-peptide response to a certain glucose level gets weaker with longer diabetes duration (Figure 5a). Diabetes duration as a quadratic term also improved the model, indicating that the decline in DBS C-peptide was not linear over time (Model 3). Finally, we tested whether age at diagnosis affected the slope of decline of C-peptide by adding an interaction term (Model 4), and this reached borderline significance, indicating that older children had a less steep decline of C-peptide over time. In the model without interactions (Model 2), a month longer diabetes duration was associated with a 17 pmol/l decline in fasting DBS C-peptide, whereas increases of 1 mmol/l in glucose, 1 year older age at diagnosis and 100 pmol/l higher baseline plasma C-peptide were associated with 18, 17 and 61 pmol/l higher fasting DBS C-peptide, respectively.

**Figure 5a.**
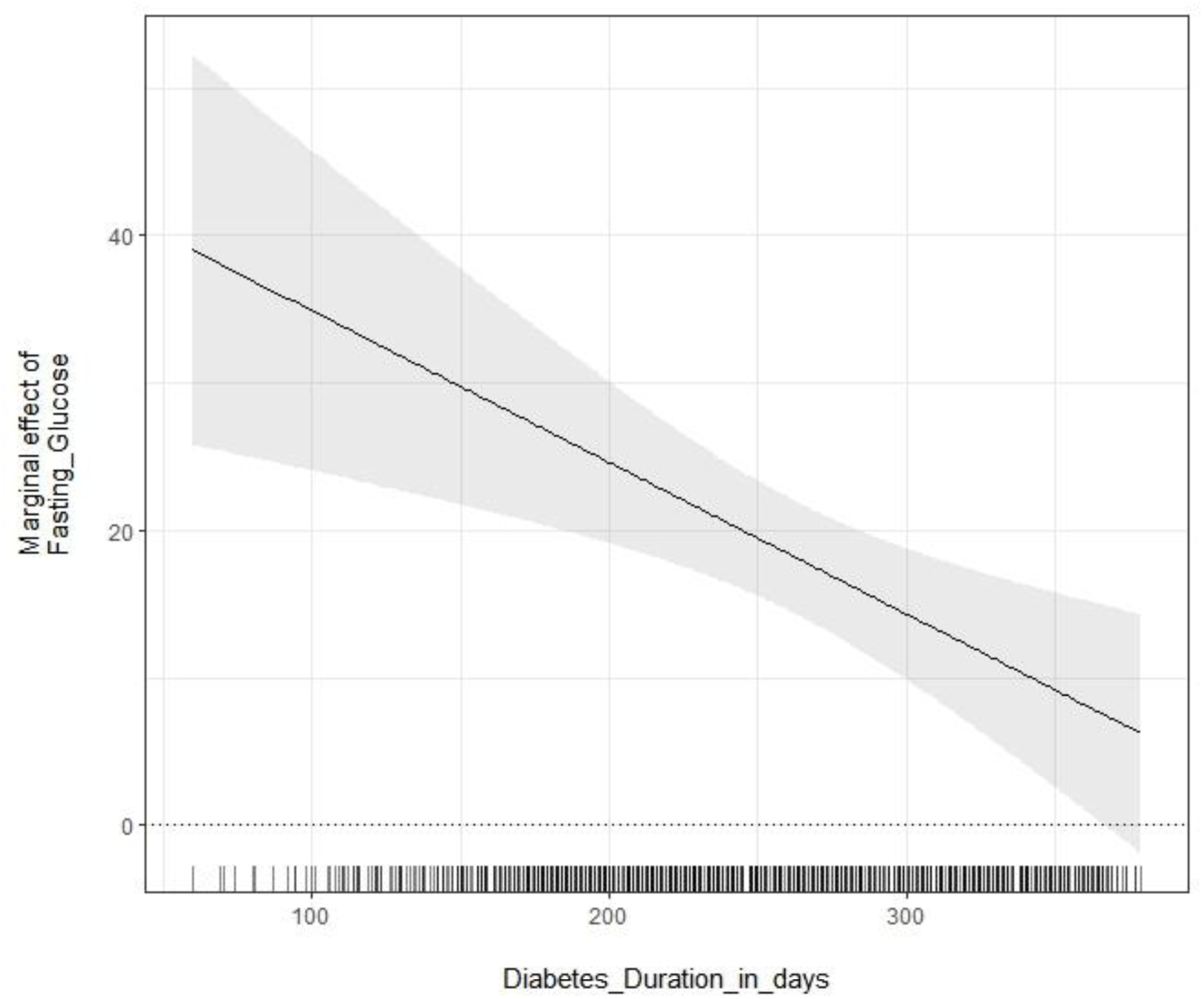
Glucose responsiveness: Marginal effect of fasting glucose on fasting DBS C-peptide levels versus diabetes duration.

### Covariates of post-prandial DBS C-peptide

Diabetes duration had a significant negative effect on post-prandial DBS C-peptide levels, whereas fasting glucose and age at diagnosis had a significant positive effect on DBS C-peptide levels (Table 2B; Model 1). Adding baseline fasting C-peptide to the model weakened the association with age at diagnosis (Model 2), likely because of confounding. Sex did not have a significant effect (data not shown). As in the fasting DBS C-peptide analyses, glucose responsiveness decreased with longer diabetes duration (Figure 5b) and the decline in DBS post-prandial C-peptide over time appeared to be non-linear (Model 3). Age at diagnosis did not affect the slope of decline of post-prandial C-peptide over time (Model 4). In the model without interactions (Model 2), a month longer diabetes duration was associated with a 42 pmol/l decline in post-prandial DBS C-peptide, whereas increases of 1 mmol/l in glucose, 1 year older age at diagnosis and 100 pmol/l higher baseline plasma C-peptide were associated with 15, 34 and 120 pmol/l higher post-prandial DBS C-peptide levels, respectively.

**Figure 5b.**
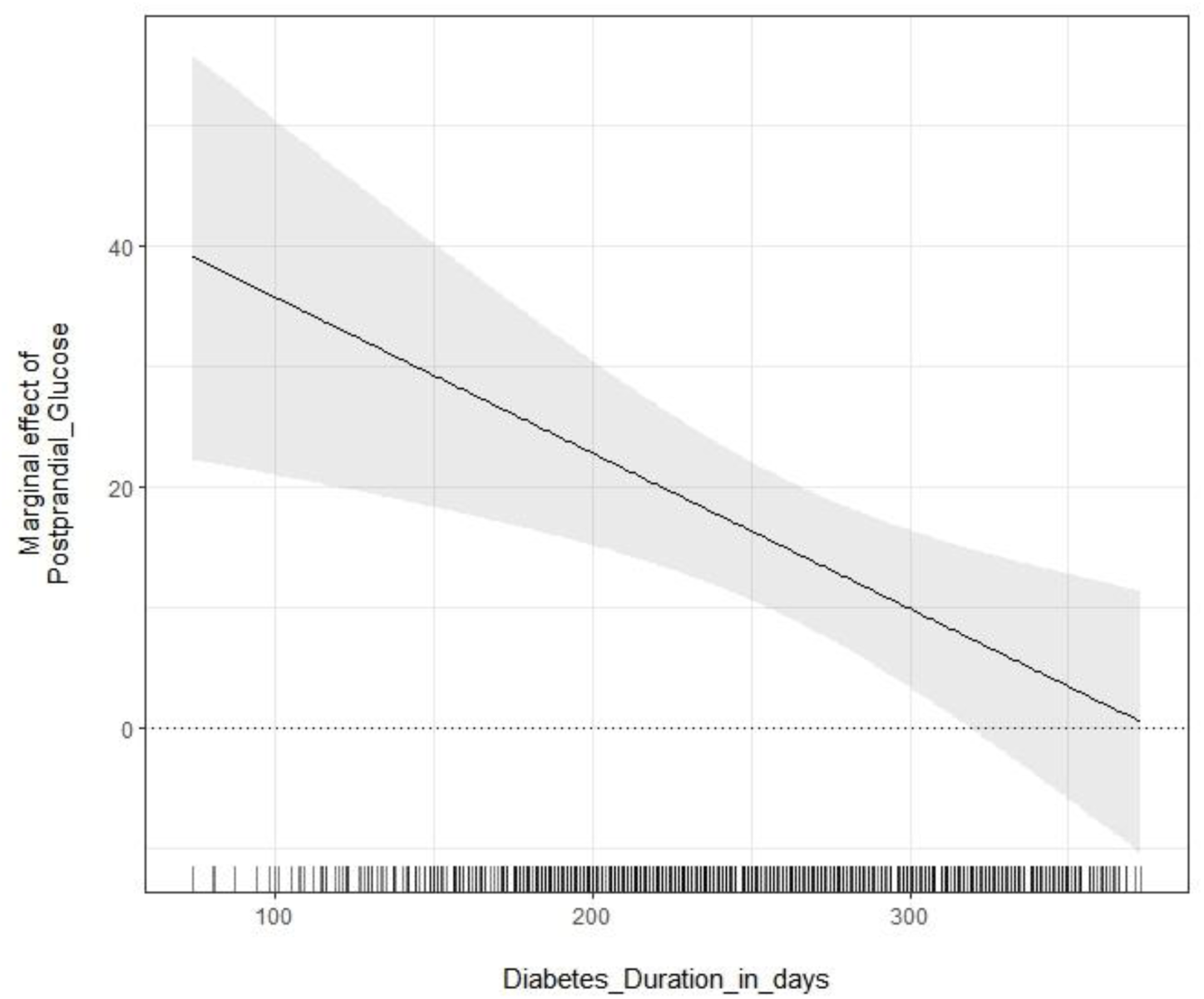
Glucose responsiveness: Marginal effect of post-prandial glucose on post-prandial DBS C-peptide levels versus diabetes duration.

**Table 2B.**
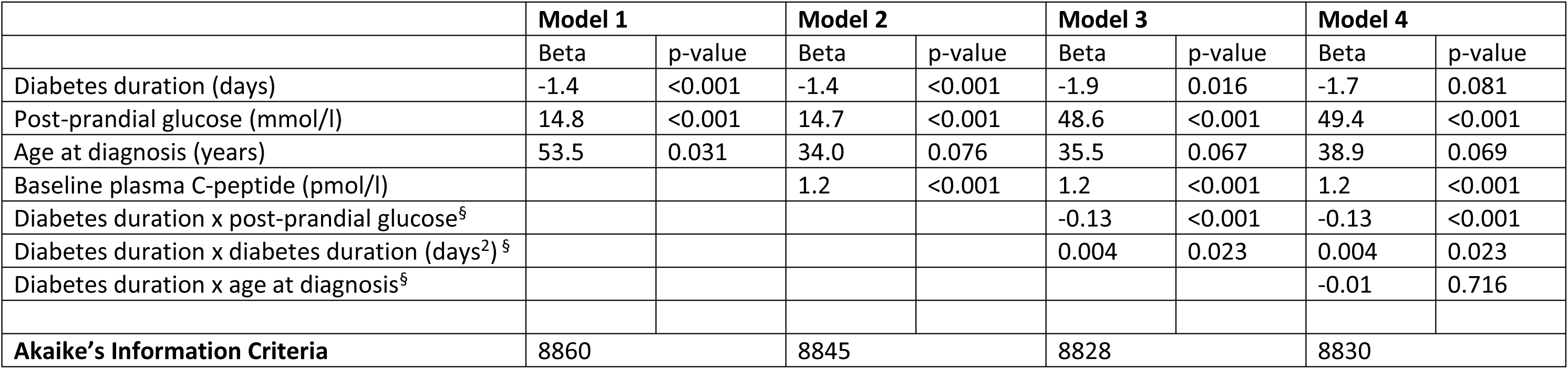
Mixed model analyses of variance: post-prandial DBS C-peptide

### Covariates of DBS C-peptide increment following breakfast

Diabetes duration and fasting DBS C-peptide had a significant negative effect on the C-peptide response to breakfast, and the glucose increment after breakfast had a significant positive effect on the C-peptide response (Table 2C; Model 1). Sex and age at diagnosis were not significant (data not shown). By adding an interaction term, we showed that with longer duration of diabetes there was a lower C-peptide response to a unit glucose increment (Model 2; Figure 5c). In the model without interactions (Model 1), a month longer duration of diabetes was associated with a 30 pmol/l decline in the DBS C-peptide increment, whereas an increase of 1 mmol/l in the delta glucose was associated with a 16 pmol/l increase in the DBS C-peptide increment, corrected for fasting DBS C-peptide levels.

**Figure 5c.**
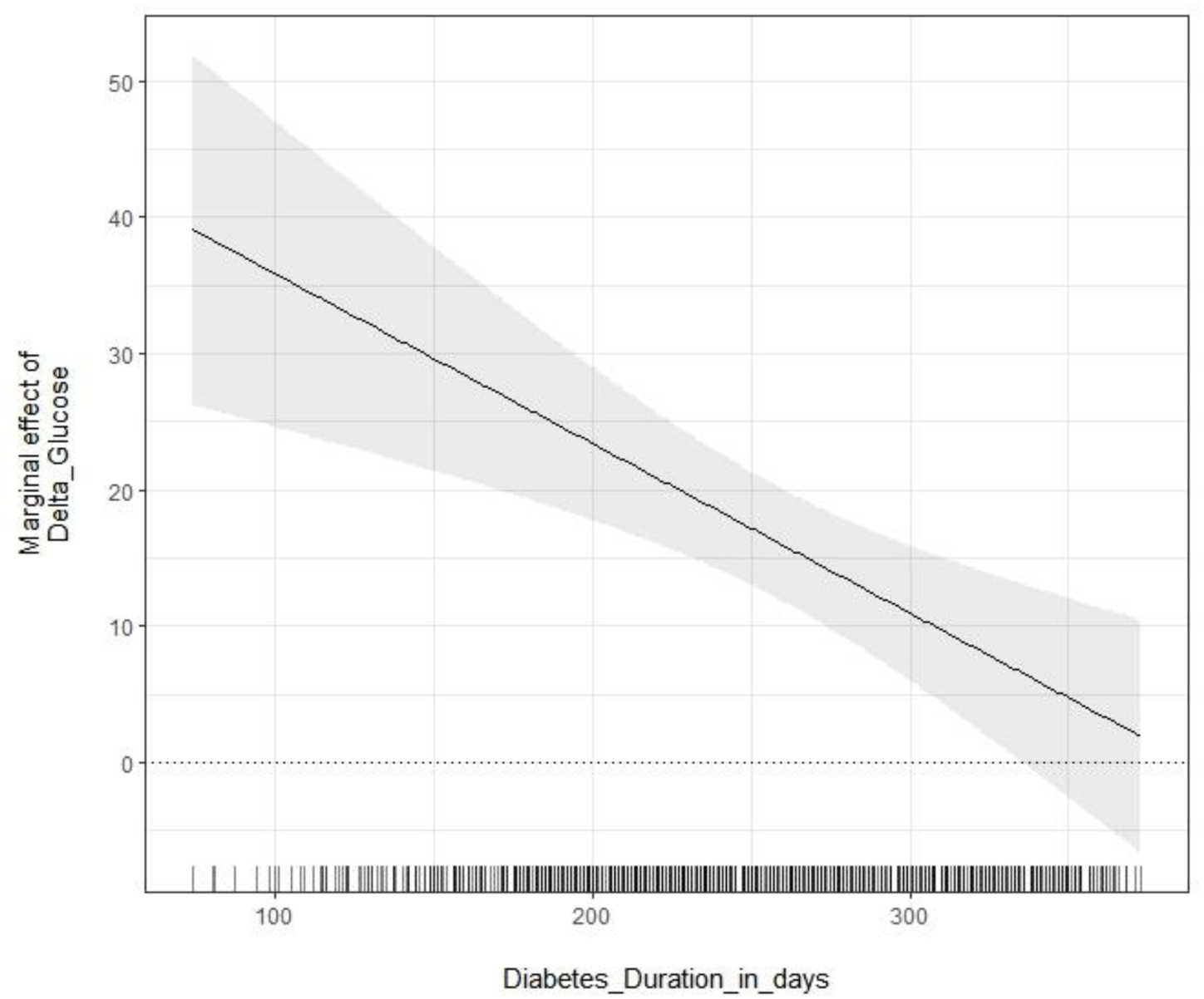
Glucose responsiveness: Marginal effect of delta glucose on DBS C-peptide increment versus diabetes duration.

**Table 2C.** Mixed model analyses of variance: DBS C-peptide increment

|  | **Model 1** |  | **Model 2** |  |
| --- | --- | --- | --- | --- |
|  | Beta | p-value | Beta | p-value |
| Diabetes duration (days) | -1.0 | <0.001 | -0.69 | <0.001 |
| Delta glucose (mmol/l) | 16.3 | <0.001 | 48.3 | <0.001 |
| Fasting DBS C-peptide (pmol/l) | -0.44 | <0.001 | -0.45 | <0.001 |
| Diabetes duration x delta glucose<sup>§</sup> |  |  | -0.12 | <0.001 |
| **Akaike's Information Criteria** | 8709 |  | 8694 |  |

<sup>§</sup>Interaction terms

## Discussion

In this study we developed an approach to measure C-peptide in DBS in children and adolescents with a recently diagnosed T1D and compared it to the MMTT. The DBS method showed a strong correlation with the plasma C-peptide method and the Bland-Altman plot indicated good performance of the DBS assay. The method proved feasible in the home setting, as indicated by the median number (24) of received DBS cards per participant. In addition, this method permitted frequent assessment of C-peptide, which allows accounting for biological variation and facilitates short-term evaluations of possible interventions.

C-peptide has been measured in DBS before and shown to remain stable on filter paper cards for 6 months (18). However, the lower limit of detection of the previous assay (18) was 440 pmol/l compared to 50 pmol/l in the current study. This increased sensitivity is important since residual concentrations of ≥200 pmol/l in T1D patients protect against diabetes complications (10,12,19).

The frequent assessment of C-peptide levels in our participants highlighted the considerable biological variation in C-peptide levels in children and adolescents with T1D. A significant proportion of this variation could be explained by the concurrent glucose levels, with higher glucose levels associated with higher C-peptide levels. In studies using the MMTT to evaluate ß-cell function, the glucose excursion during a MMTT is normally not taken into account, but our data suggest that this has considerable impact on measured C-peptide levels. Future studies evaluating potential biomarkers of an individual’s change in ß-cell function over time could adjust for concurrent glucose levels.

By investigating which covariates had an effect on DBS C-peptide levels, we demonstrated, as expected, that longer diabetes duration was associated with lower DBS C-peptide levels. The decline in C-peptide over time was not linear but curved, which has been described before in a study using MMTTs, but in that study the slope of decline in ß-cell function only changed after 12 months diabetes duration (5). Concurrently measured glucose levels had a positive impact on C-peptide levels. In addition, we demonstrated that this positive effect of glucose on C-peptide levels decreases with longer diabetes duration. This was apparent in all three models, using fasting DBS C-peptide, post-prandial DBS C-peptide or the DBS C-peptide increment as the dependent variable. This indicates that the ability of the pancreas to secrete more insulin in response to higher glucose levels, also called ‘glucose responsiveness’ or ‘glucose sensitivity’ (20), decreases with longer duration of diabetes. A decrease in this ‘glucose sensitivity’ was found previously to be an early and strong predictor of progression to T1D in at risk family members (20). The DBS method has not been evaluated yet in at risk individuals, but the fact that changes in glucose responsiveness could be picked up is encouraging.

As expected, greater age at diagnosis was associated with both higher fasting and post-prandial C-peptide levels. We also demonstrated that younger children had a steeper decline of fasting C-peptide levels, in line with the results of a large study including 3929 patients from seven European registries, that demonstrated a more rapid decline in fasting plasma C-peptide in those with earlier onset of T1D (21). Ludvigsson et al. reported a greater difference between random, non-fasting C-peptide at diagnosis and at one year from diagnosis for those with a younger age at diagnosis, suggesting that younger children have a more rapid decline of their ß-cell function (7). The decline in C-peptide seemed particularly pronounced in children <5 years at diagnosis (7). Greenbaum et al. also reported a slower decline of ß-cell function in subjects older than 21 years of age compared to younger patients, but a similar rate of decline in individuals aged 7-21 (5).

A limitation of our study is the fact that we did not ask the participants to withhold insulin in the home setting, and this may have influenced the amplitude of the prandial C-peptide response. We anticipated that omitting insulin at home would be too disruptive and might negatively affect adherence to the study. In retrospect this was unnecessary as indicated by the high number of collected DBS cards. Previous research has shown that the peak C-peptide in a MMTT with concurrent insulin administration, albeit lower, was still highly correlated to the peak in a MMTT without insulin (22). Another limitation is that the participants did not have the exact same stimulus for C-peptide secretion at home, as their breakfasts may have differed in fat and protein content, although we ensured age-banded, comparable amounts of carbohydrates in the breakfast across all participants and the weekly breakfast for each participant was the same.

Our study highlights the heterogeneous nature of changes in ß-cell function in children and adolescents with T1D. Some participants showed a dramatic decline in ß-cell function within the first year from diagnosis, whereas others maintained their ß-cell function relatively well. There were only two participants whose ß-cell function became undetectable during the study. The identification of biomarkers that can predict the course of ß-cell function within the first year from diagnosis will be of major importance, as it would inform which participants are likely to benefit most from interventions. In a previous study sCD25, an established marker of immune activation and inflammation, was found to be elevated in T1D patients as compared to controls, and was also negatively associated with C-peptide levels (23). This marker and other markers of immune activity such as C-reactive protein could also be measured from DBS. Simultaneous measurement of metabolic and immune status could be particularly valuable in future attempts to stratify patients, and in the frequent monitoring of the effects of potential therapeutics.

There’s increasing interest in interventions to preserve or even restore residual ß-cell function in T1D patients (9). Some drugs approved for treatment of type 2 diabetes have been investigated in paediatric T1D, such as thiazolidinediones (24) and glucagon-like peptide 1 receptor agonists (25), and there’s a growing number of immunotherapy studies (26–31).

Some T1D trials have reported some, if temporary, beneficial effects on ß-cell function (26,27,29,31), although this only led to reductions in insulin requirements in a minority (29, 31). Others reported beneficial effects in adults, but not in children (28) and with the opposite trend for a promising approach, inhibition of the T cell receptor CD3 (32). T1D is a heterogeneous disease and it’s likely that a more tailored, stratified approach is required to improve and optimise treatment success. Such adaptive and dynamic treatment approaches are on their way (33), and a straightforward, less labour-intensive method permitting more frequent assessment of ß-cell function may aid in the evaluation of interventions. Frequent assessment of C-peptide via DBS enables us to calculate a slope reflecting the change in ß-cell function over time. The characterisation of an individual’s slope following diagnosis may help in selecting participants which are likely to benefit most from intervention trials and analysing changes in this slope may provide useful in the short term evaluation of promising interventions. Finally, the DBS method may be useful in monitoring ß-cell function in auto-antibody positive at risk individuals, a feature of T1D research receiving increasing attention (34).

## Acknowledgements

We thank all participants and parents for participating in the study. We greatly acknowledge Katrin Mooslehner, Radka Platte, Karen Whitehead and Di Wingate for their dedicated help with sample processing; Kayleigh Aston, Kimberley Dale, Andy Kempa, Clare Megson, Monica Mitchell and Criona O’Brien, research nurses, the staff at the National Institute for Health Research (NIHR)/Wellcome Trust Clinical Research Facility Cambridge and NIHR/Wellcome Trust Clinical Research Facility, Birmingham Children’s Hospital, for recruitment of participants, assistance and help with data collection; and Roman Hovorka and Ken Ong for help with the statistical analyses.

---

**Supplementary Figure 1.**
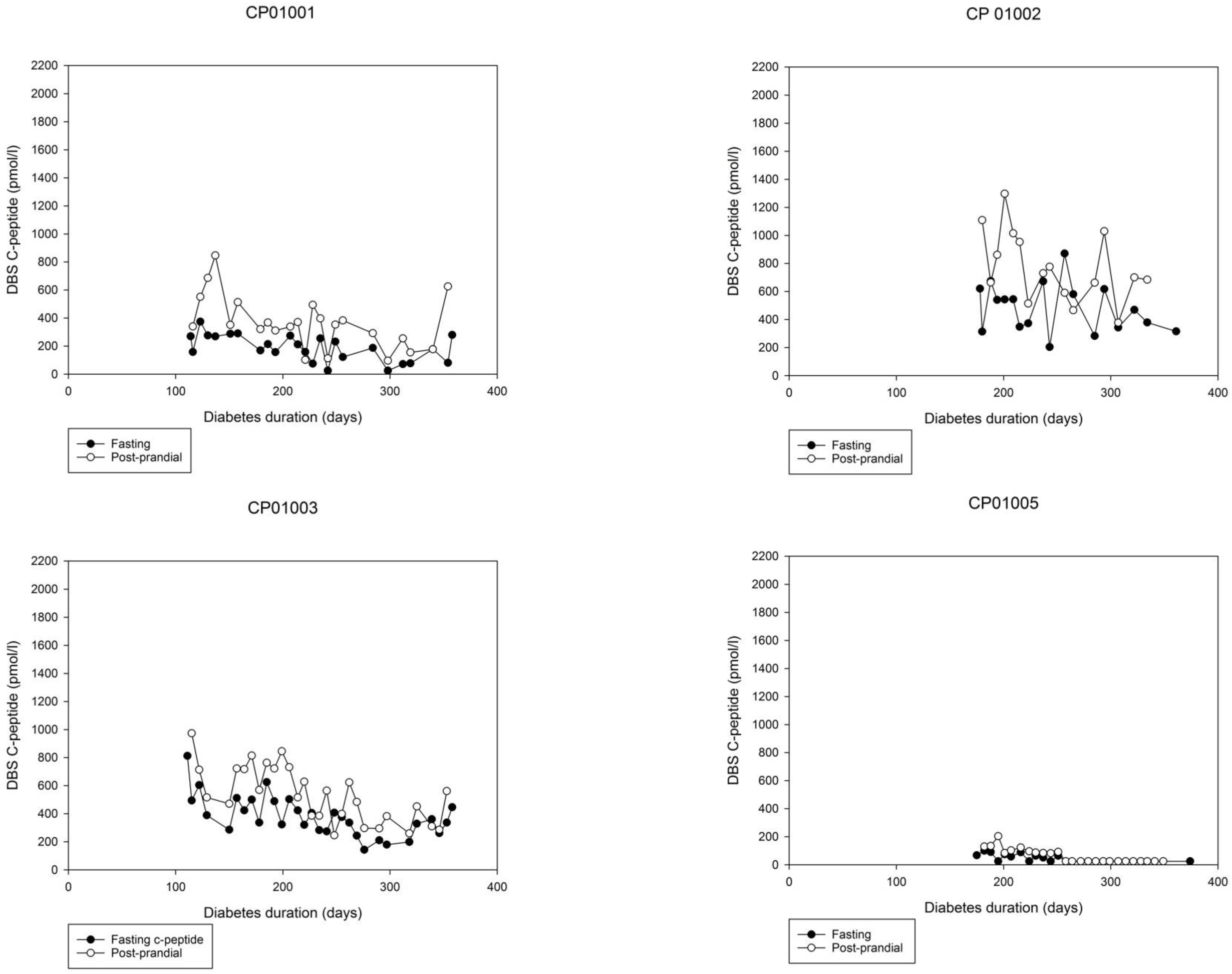
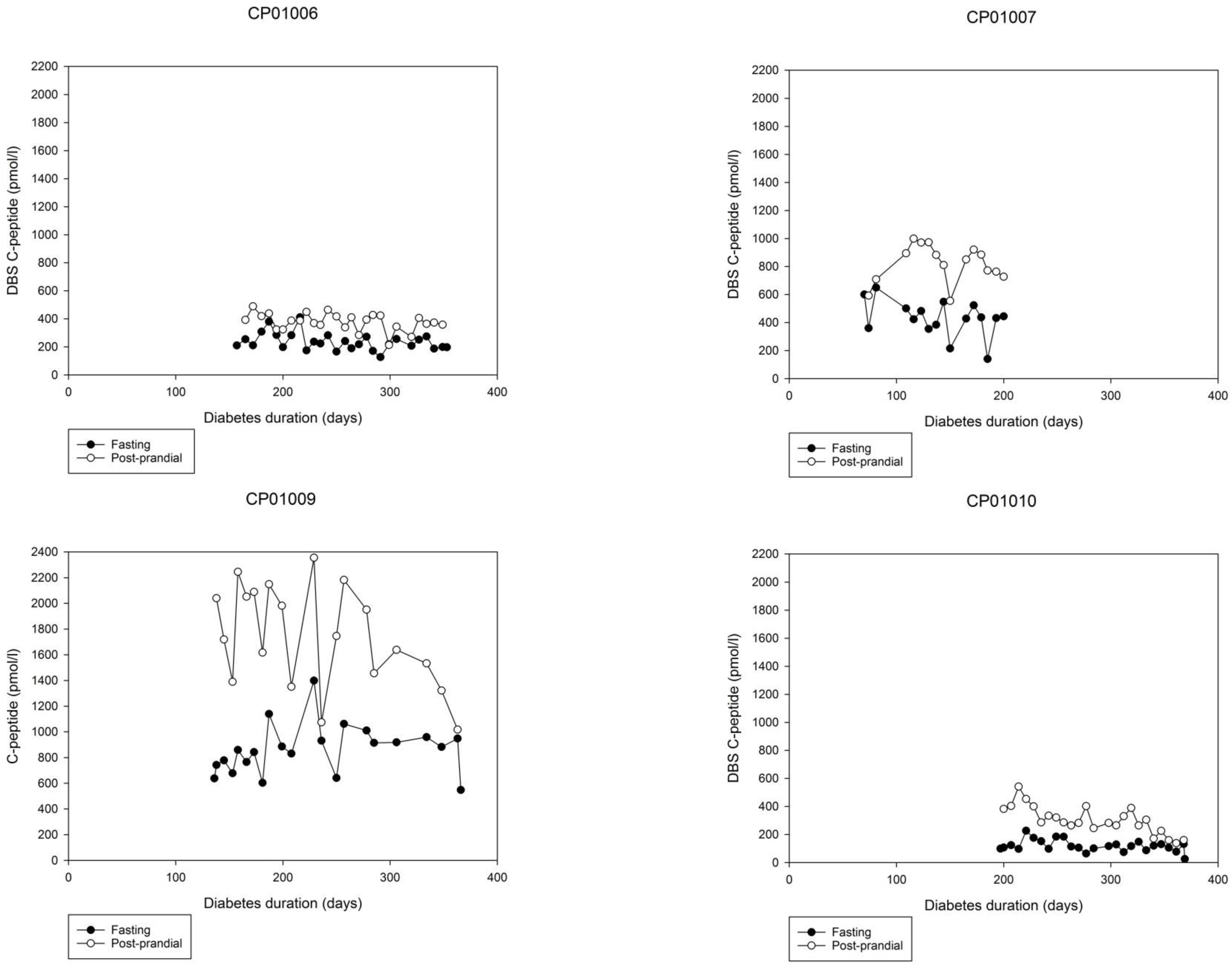
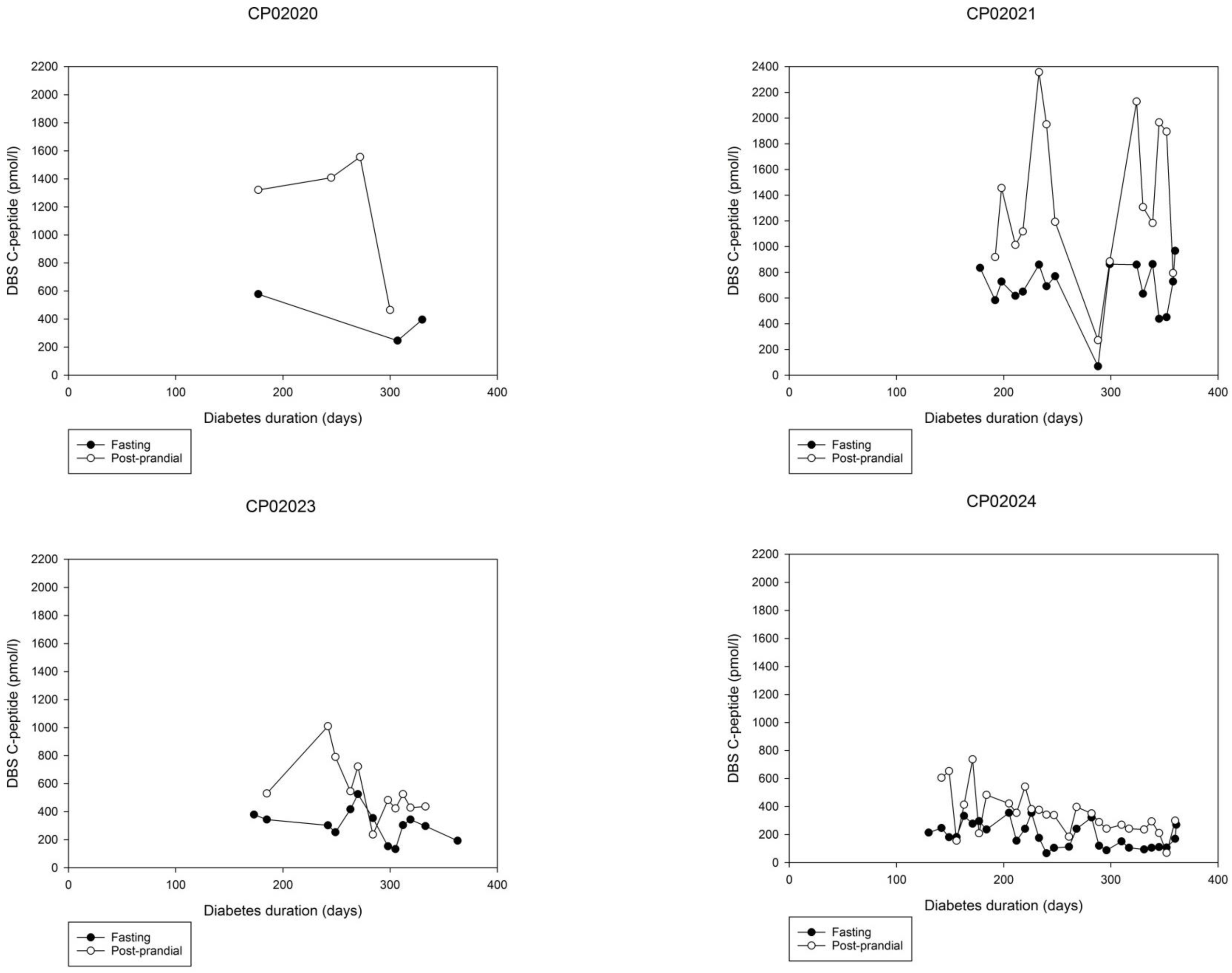
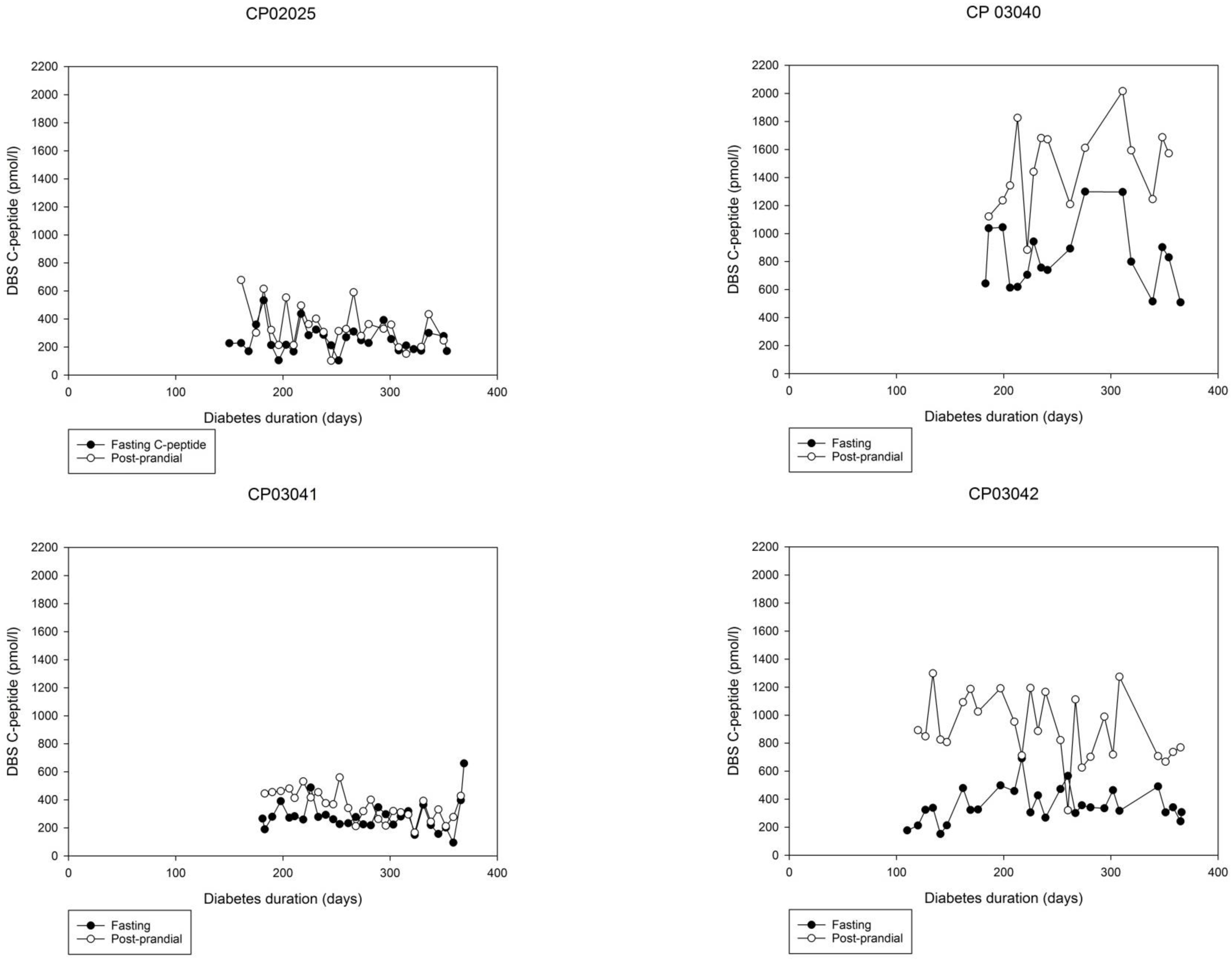
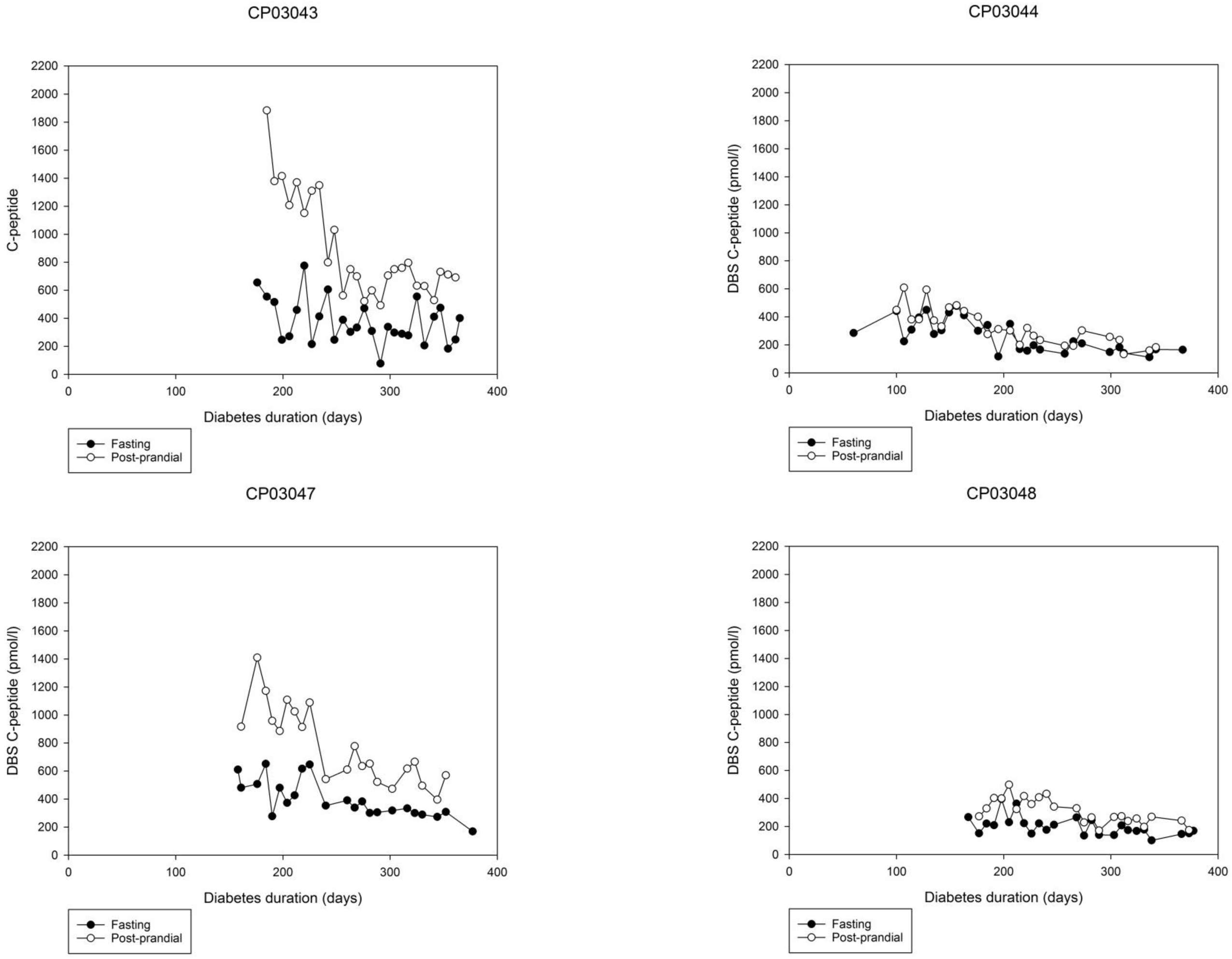
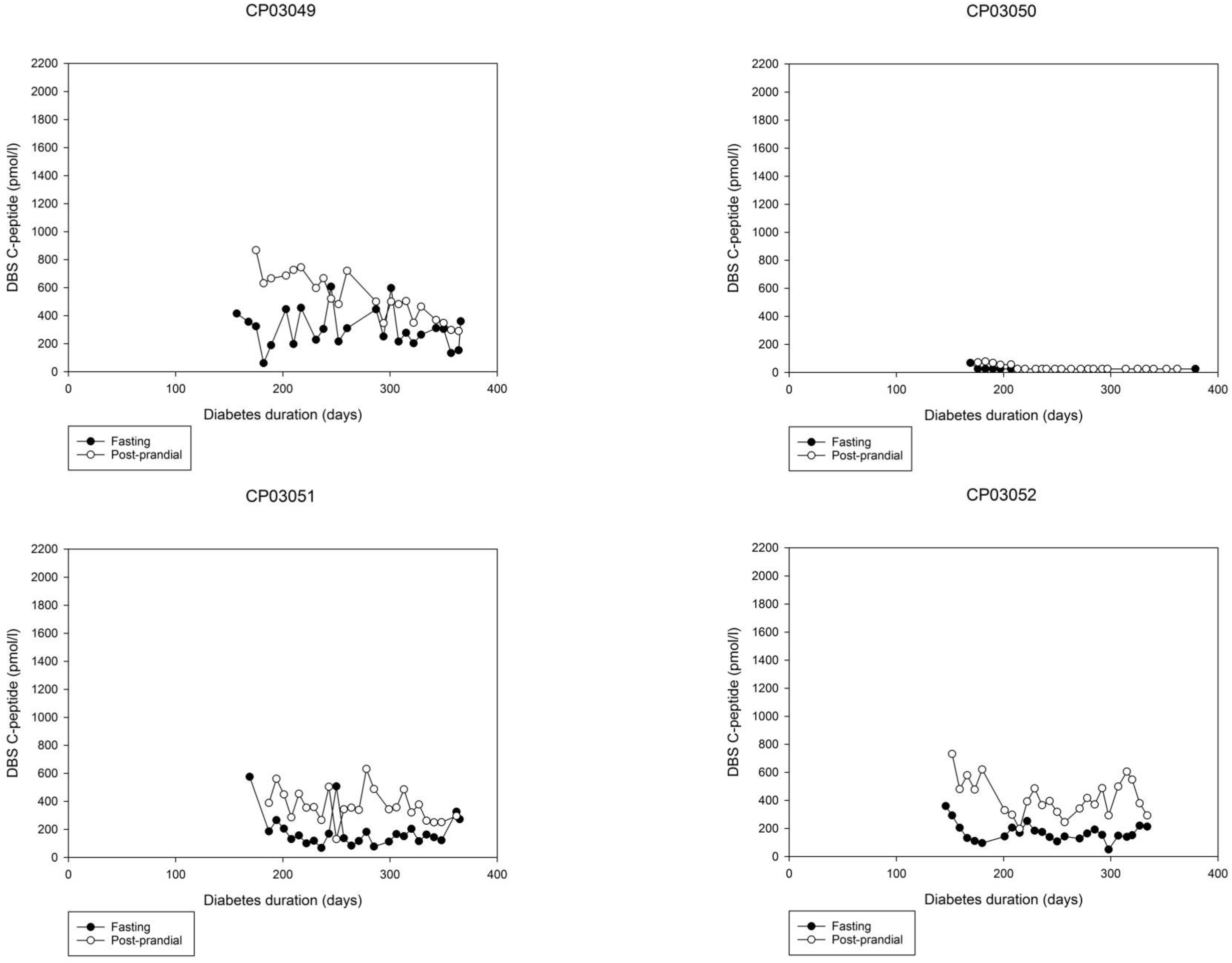
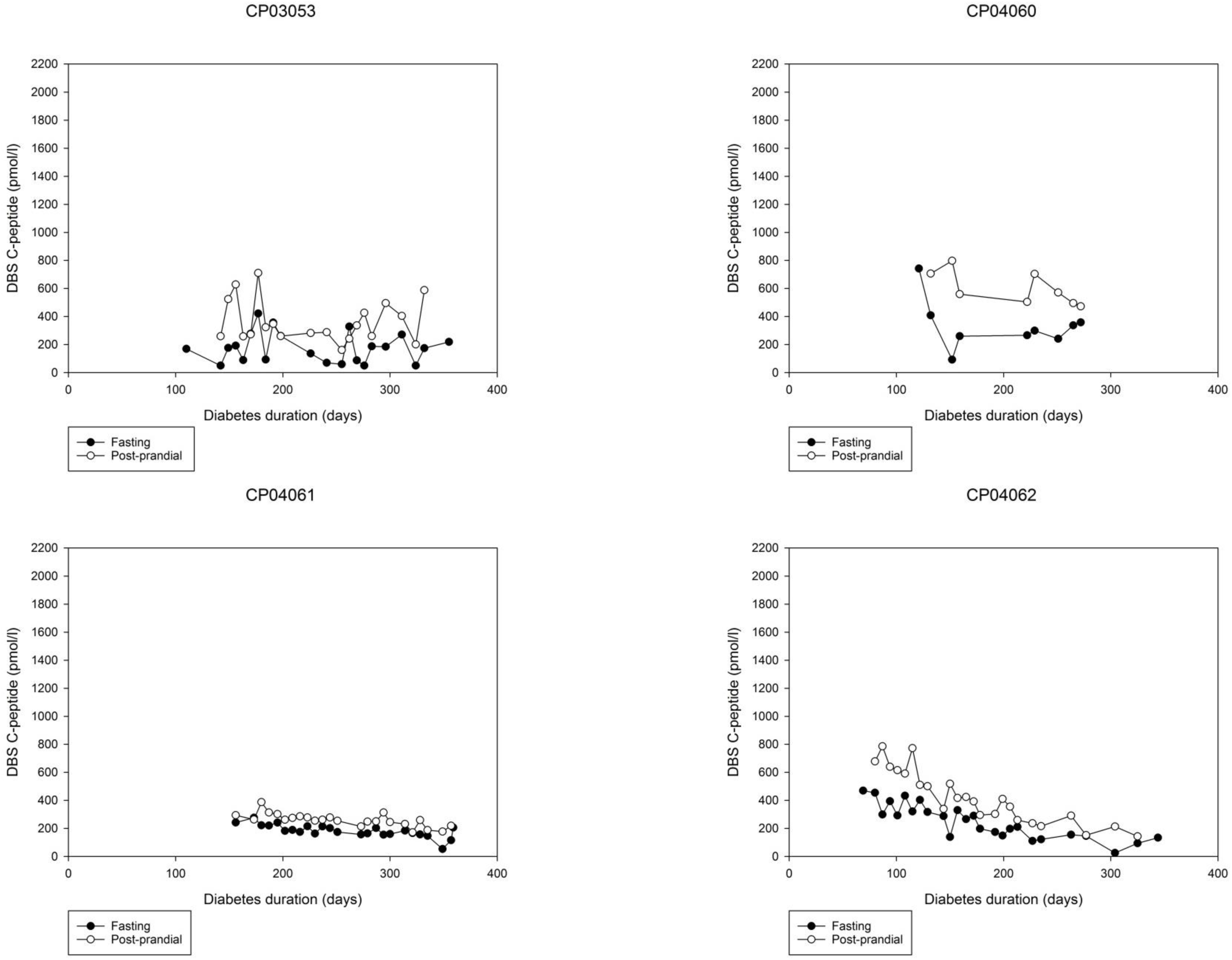
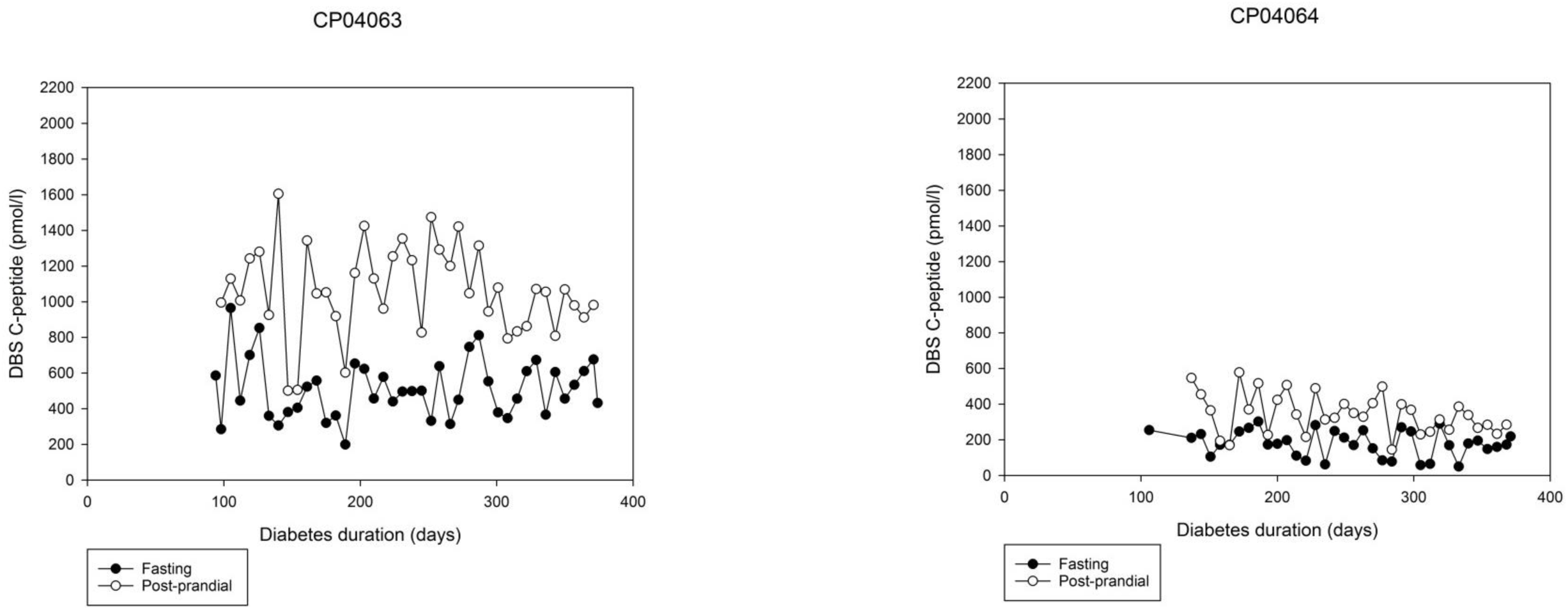
Individual participants’ course of fasting and post-prandial DBS C-peptide vs. diabetes duration

